# Rapid quantification of short-chain fatty acids by Fourier-transform infrared spectroscopy for microbiota quality assessment

**DOI:** 10.1101/2025.11.25.690441

**Authors:** Anna Sayol-Altarriba, Andrea Aira, Eva Martín-López, Anna Villasante, Rosa Albarracín, Joana Faneca, Cristina Pitart, Ignasi Roca, Gregori Casals, Santiago Marco, José Luis Villanueva-Cañas, Climent Casals-Pascual

## Abstract

Short-chain fatty acids (SCFAs) are bacterial metabolites with crucial roles in host homeostasis and immune system modulation. Given their benefits, they have been proposed as markers of a healthy microbiota. Their quantification, however, is time-consuming, expensive and requires specialized personal and equipment. Thus, their use for quality assessment of stool samples in clinical contexts is limited. In this study we explored the use of Fourier-transform infrared (FT-IR) spectroscopy as an alternative method for metabolomic quality assessment of stool samples. FT-IR identifies the chemical bonds present in SCFAs and allows clear discrimination between healthy and dysbiotic stool samples (caused by *Clostridioides difficile* infection) through Principal Component Analysis (PCA). Overall, FT-IR provides a rapid, cost-effective and simple method for stool samples quality assessment. Combined with the widespread availability of this technology in most hospitals, these advantages highlight its potential application for routine implementation in clinical laboratories.

## INTRODUCTION

Short-chain fatty acids (SCFAs) are key metabolic by-products synthesized by gut bacteria. They are produced in the colon through anaerobic fermentation, mainly from dietary fibres non-digestible by the host, but also from amino acids that have not been absorbed and reach the colon [1]. The main SCFAs in terms of abundance and functionality are acetic, propionic and butyric acids, mostly found in their anionic form in the human colon (acetate, propionate and butyrate) [2].

There is growing evidence of the prominent role of these metabolites in both gut and systemic health, where they contribute to the maintenance of gut integrity and homeostasis [3],serve as energy sources for colonocytes (mainly butyrate) and for other cells (acetate and propionate) [4], are involved in the systemic energy metabolism [5], show anti-inflammatory, antimicrobial and antitumorigenic activities [6], and participate in the modulation of the immune system [6]. By playing such prominent roles, SCFAs have implications in several diseases such as gastrointestinal disorders [7], metabolic diseases [8,9], and even certain cancers [10]. All these implications highlight the need for a reliable quantitative and qualitative detection methods in clinical laboratories, which are crucial for clarifying their specific roles in several pathologies and for implementing their use as functional biomarkers. This is of special relevance for the implementation of faecal microbiota transplantation (FMT) beyond recurrent *Clostridioides difficile* infections (CDI).

Currently, there are several methods available for SCFAs quantification. The most common method is gas chromatography coupled with mass spectrometry (GC/MS), which provides high-accuracy and sensitivity but requires complex protocols, technical expertise, specialized instruments, long running times, high costs of supplies and hazardous chemicals [11,12]. Other alternatives are high-performance liquid chromatography (HPLC), nuclear magnetic resonance (NMR) and capillary electrophoresis (CE) [13–15]. While each technique offers specific advantages and limitations, but there is an overall unmet need for a fast, cost-effective and simple method for the quality assessment SCFAs in stool samples. More recently, paper-loaded direct analysis in real time mass spectrometry (pDART-MS) has been proposed as a novel method to quantify SCFAs [16]. While this method offers a quick and high-throughput analysis, it still requires highly specialized instrumentation, which limits its potential for routine use in clinical laboratories.

In this context, infrared spectroscopy offers a promising alternative methodology. In particular, Fourier-transform infrared (FT-IR) spectroscopy has recently gained attention in clinical microbiology for real-time outbreak detection and strain-typing of bacteria [17,18]. In other fields such as the food industry, IR-based technologies have been applied for many years as quality control tools; for instance, to quantify SCFAs contributing to cheese flavour or to detect fatty acids (including SCFAs) in bovine milk [11,19]. In human samples, FT-IR has also been tested for quantifying SCFAs in breast milk, given their importance in infant health; Near-IR spectroscopy has similarly been explored as a novel method to quantify SCFAs, although with limited performance [20,21].

In FT-IR, the interaction of IR radiation with molecules in a sample is used to measure the vibrational frequencies of their chemical bonds and functional groups through changes in absorption. The intensity of absorbed light is measured across different wavenumbers in the mid-IR spectrum (4,000 to 400 cm^-1^), producing a characteristic spectrum of the sample. Once processed, this spectrum can be used to evaluate functional groups and molecules, both in a qualitative and quantitative manner [17,22]. Compared with other available methods, FT-IR is cost-effective, has minimal operational costs, allows analysis of sample within minutes, provides functional information, and does not require highly trained personnel. All these features make it a promising method for stool sample quality assessment and support its potential incorporation into routine protocols.

In this study we have evaluated FT-IR as a novel methodology for assessing the metabolomic quality of stool samples. We have compared stools from healthy donors, obtained from the Stool Bank at Hospital Clínic de Barcelona, with those from dysbiotic patients with CDI. All samples were also analysed by GC/MS, the current gold-standard technique for SCFAs quantification. Overall, our results suggest that FT-IR could facilitate the robust and rapid quality assessment of stool samples for routine use in clinical laboratories.

## MATERIALS AND METHODS

### Sample selection and conservation

Samples from healthy donors and patients with dysbiosis caused by CDI were collected and conserved following the criteria described in previous studies [23]. Briefly, samples from healthy donors (n = 115) were obtained from surplus samples at the Stool Bank of Hospital Clínic de Barcelona (2021 – 2022), and stool samples from patients with dysbiosis caused by CDI (n = 40) were collected from stool sample leftovers at the Department of Clinical Microbiology of Hospital Clínic de Barcelona. All samples used for the study were anonymized using an alphanumeric code. The study was approved by the Ethics Committee for Research with medicines (CEIm) of Hospital Clínic de Barcelona (Ref. HCB/2024/0239). Informed consent from CDI patients was waived by the ethics committee as the samples used were from surplus material from the Clinical Microbiology Department and no clinical or personal data were accessed.

### SCFAs quantification

SCFAs were analysed by gas chromatography coupled with mass spectrometry (GC/MS) following our previously described protocol, including the normalisation of SCFAs levels by the bacterial count of the original stool sample [23].

Briefly, stool samples were acidified with hydrochloric acid (HCl) in water to a final concentration of 100 mg of stool/mL and centrifuged at 13,000 rpm for 3 min. Half of the supernatant was transferred into a fresh tube and stored at - 20 ºC until SCFAs quantification by GC/MS (∼ 2 months). The remaining part of the supernatant was stored at - 20 ºC until used for the FT-IR spectroscopy analysis (∼ 1 month, see below).

For GC/MS quantification, samples were further centrifuged, extracted using anhydrous diethyl ether, derivatized with N,O-bis(trimethylsilyl)trifluoroacetamide (BSTFA) and trimethylchlorosilane (TMCS) (99:1), and injected into a Shimadzu GCMS-QP2010 Ultra instrument.

### FT-IR measurement

For FT-IR spectroscopy assessment, tubes with the acidified supernatant of stool samples were thawed at room temperature and 15 µL of the supernatant was placed into a 96-well FT-IR silicon plate and allowed to dry. Five technical replicates per sample were used (except for two samples, which had four technical replicates). An IR Biotyper® instrument (Bruker Daltonics GmbH & Co KG, Bremen, Germany) was used for all the FT-IR measurements. In each run, Bruker Infrared Test Standards 1 and 2 (ITRS 1/ITRS 2) were included for quality controls. Spectra (3,996 – 500 cm^-1^) were acquired using OPUS version 8.2.28 software and IR Biotyper® version 4.0 software (Bruker Daltonics GmbH & Co KG, Bremen, Germany) with default settings (32 scans per technical replicate; spectral resolution, 6 cm^-1^; apodization function, Blackman Harrys 3-term; zero-filling factor, 4).

To ensure that the signal captured by the FT-IR spectroscopy instrument was specific to the metabolic component of our samples, we performed two tests to optimize the signal-to-noise ratio. First, we ran a negative background control using the acidified water used for the supernatant preparation. Second, we tested four random samples mixed in pairs in equal proportion, to minimize random signal pick-up. In both cases, the samples were prepared and scanned with the FT-IR instrument using the same volume, replicates and parameters for the spectra acquisition previously described.

### Spectra processing and statistical analysis

The spectra processing and statistical analysis were performed using R through RStudio software version 4.3.1 (RStudio: Integrated Development Environment for R, Boston, MA, USA). Data from OPUS binary files were extracted using the opusreader2 R package [24], and processed using prospectr R package [25]. FT-IR full replicate spectrums of each sample (3,996 - 500 cm^-1^) were averaged, corrected with: 1) the Standard Normal Variate (SNV), 2) baseline removal and 3) the Savitzky-Golay smoothing and second derivative calculation (derivative order, m = 2; polynomial order, p = 3; window size, w = 9). After processing the spectra, one CDI sample was discarded for the subsequent analysis as it was considered an outlier (final CDI n = 39). As the calculation of the second derivative enhances the resolution and the separation of overlapping peaks, the resulting total processed spectra (3,992 - 504 cm^-1^) were subsequently used for the principal component analysis (PCA) using the FactoMineR [26] and the factoextra R packages [27].

## RESULTS

### Classification of healthy and CDI dysbiotic stool samples using characterized FT-IR absorbance

The SNV corrected spectra after baseline removal from both groups (healthy donors and dysbiotic patients with CDI) were averaged respectively and analysed according to groups and bonds found in SCFAs (acetic, propionic and butyric acids) (Figure 1C, butyric acid is shown as a representative of SCFAs). The general absorbance of samples from CDI dysbiotic samples was lower than the absorbance in samples of healthy donors. The spectra showed a broad absorbance band between 3,600 and 3,100 cm^-1^, attributable to the O-H stretch vibration of the hydroxyl group of carboxylic acids; asymmetric (out-of-phase) and symmetric (in-phase) stretching bands between 3,000 and 2,800 cm^-1^, characteristic of C-H vibrations of methyl (CH_3_) and methylene (CH_2_) groups; strong bands between 1,700 and 1,400 cm^-1^ corresponding to bending and stretching of the C=O bond of the carbonyl group of carboxylic acids and acid salts; and a weak to moderate band between 1,380 and 1,210 cm^-1^ attributable to the C-O stretch of carboxylic acids. Aliphatic groups in SCFAs can also show a deformation band of the CH_2_ bend around 1,464cm^-1^, overlapping with the C=O region (Figure 1B).

**Figure 1.**
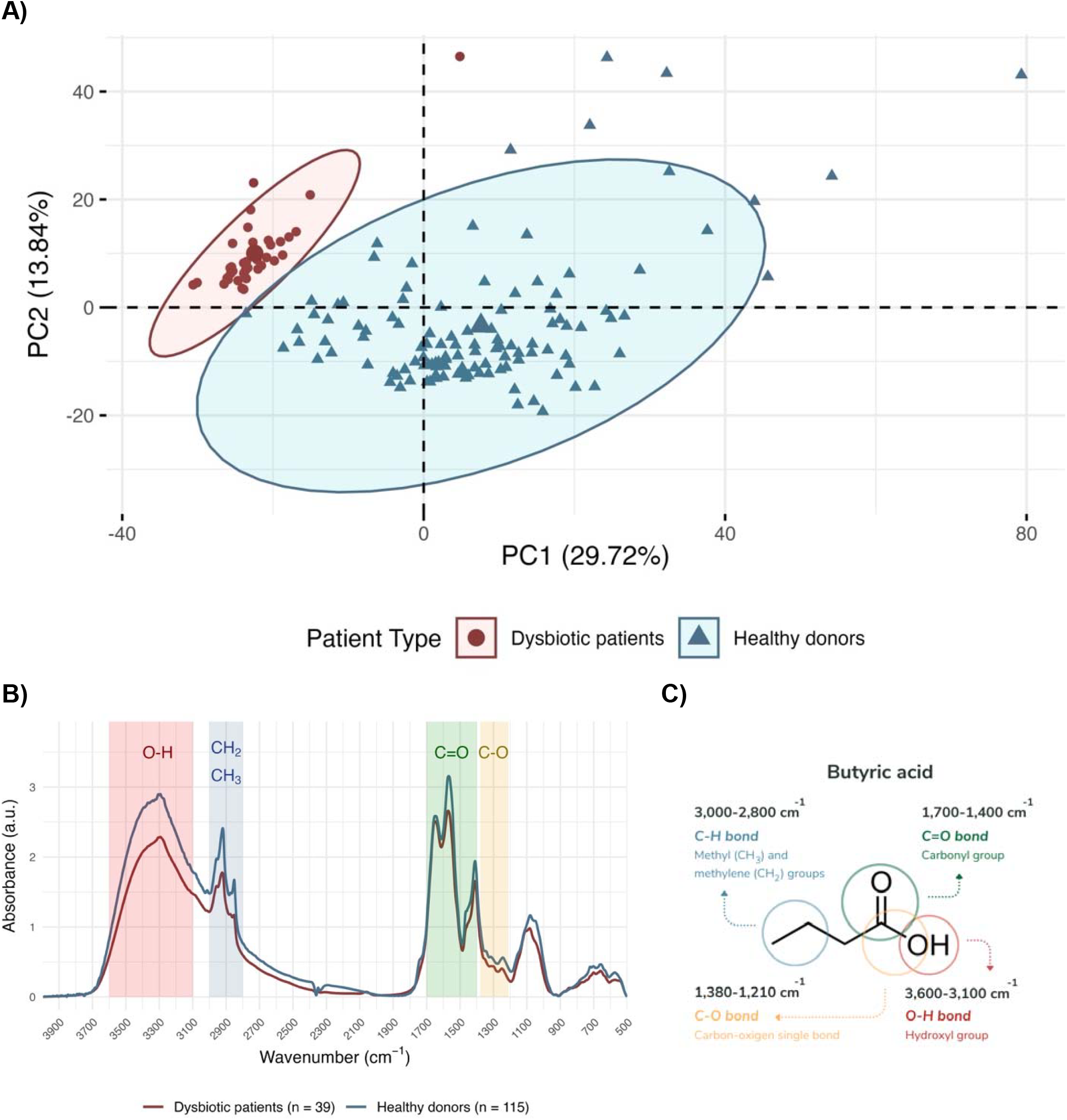
FT-IT analysis of faecal supernatants. **(A)** PCA score plot built from selected regions of the normalised FT-IR spectra (SNV correction, baseline removal, with Savitzky-Golay second derivative) of acidified stool samples from dysbiotic patients with CDI (n = 39, red dots) and healthy donors (n = 115, blue triangles). Ellipses indicate the 95% confidence interval of each group. **(B)** Averaged absorbance (a.u., arbitrary units) of mid-infrared (3,996 - 500 cm^-1^) spectra of acidified stool samples from dysbiotic patients with CDI and healthy donors, corrected with the SNV and baseline removal. Bands for specific groups and bonds are marked in the spectra: O-H band (3,600 – 3,100 cm^-1^, red), C-H band (3,000 – 2,800 cm^-1^, blue), C=O band (1,700 – 1,400 cm^-1^, green), C-O band (1,380 – 1,210 cm^-1^, yellow). Band assignments adapted from Larkin, 2011. **(C)** Chemical structure of butyric acid (CH_3_CH_2_CH_2_COOH), representative of the chemical groups and bonds present in SCFAs and their detection regions in the FT-IR spectra.

After variable autoscaling, PCA was conducted using two approaches: 1) the absorbance values from selected spectral regions (Figure 1B) in the normalised FT-IR spectra (SNV, baseline removal, and Savitzky-Golay second derivative) corresponding to chemical bonds in SCFAs (Figure 1A), and 2) the full normalised FT-IR spectra (3,992 – 504 cm^.1^) (Supplementary Figure S2). The analysis restricted to SCFA-related regions yielded a clearer separation between groups in the PCA score plot. Overall, 153 PCs were identified, of which 67 met Kaiser’s criterion (eigenvalues > 1). PC1 and PC2 accounted for 29.72% and 13.84% of the variance, respectively, explaining a combined 43.56%.

The contribution of individual variables to the PCA score plot was assessed (Supplementary Figure S3). For PC1, the 100 variables with the highest loadings corresponded mainly to absorption bands attributed to the O–H bond, as well as CH_2_ and CH_3_ groups. In contrast, the variables contributing most to PC2 were associated with CH□ and CH□ groups, with a smaller contribution from C=O bond absorptions.

### Association between FT-IR spectral signatures and SCFAs levels measured by GC/MS

All samples, including those obtained from healthy donors and CDI dysbiotic patients, were ranked according to their normalised SCFAs concentrations measured by GC/MS. Based on this ranking, we divided the dataset into deciles, and specifically focused on the extreme groups: samples within the lowest 10% (below the 10^th^ percentile) and the highest 10% (above the 90^th^ percentile) for each individual SCFAs. Using the selected FT-IR corrected spectral regions, we then performed a PCA to evaluate whether spectral profiles could discriminate between these two groups. Score plots were generated for acetate (Supplementary figures 4A and 4B), propionate (Supplementary figures 4C and 4D), and butyrate (Figure 2A and 2B). Among these metabolites, butyrate showed the clearest separation between the lowest and highest deciles, while acetate and propionate showed some level of overlap between groups.

**Figure 2.**
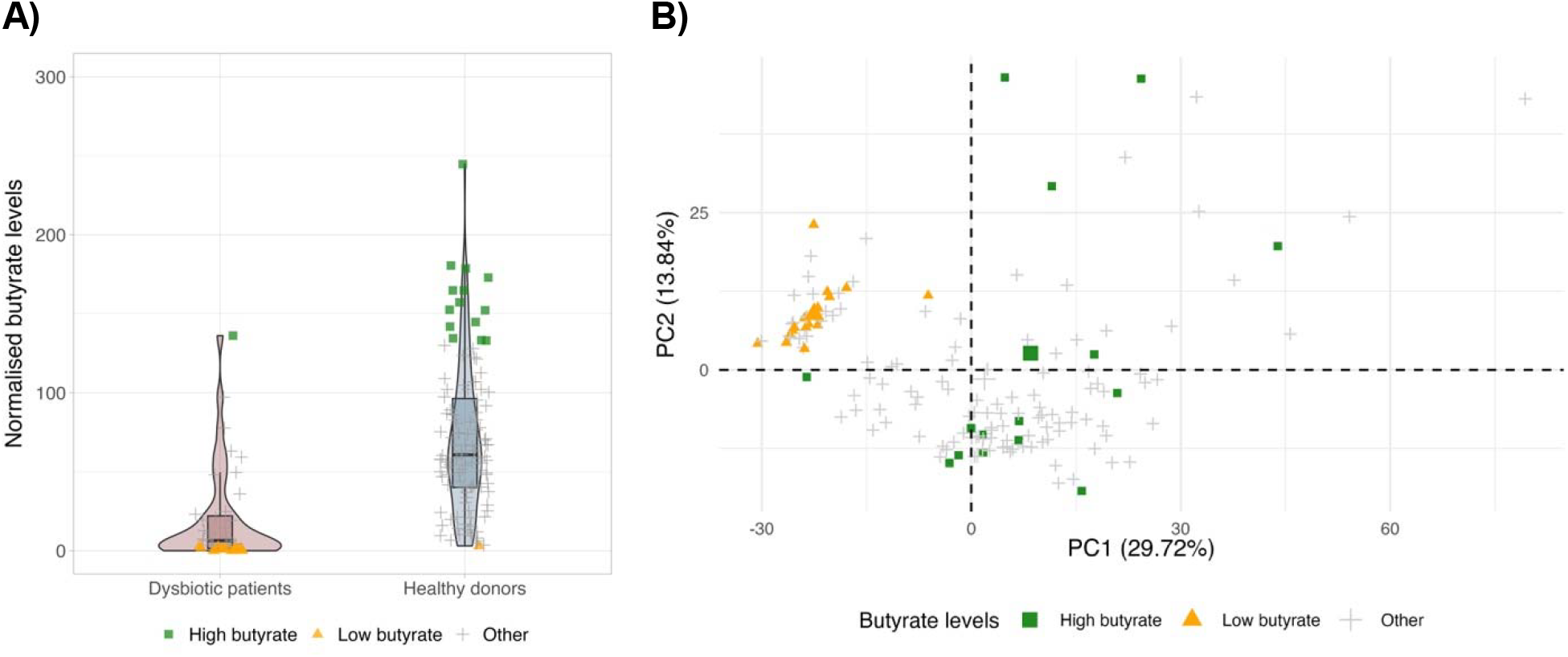
Association between FT-IR spectral signatures and SCFAs levels measured by GC/MS. **(A)** Normalised butyrate levels by bacterial count in dysbiotic patients with CDI (n = 39) (red) and healthy donors (n = 115) (blue), measured by GC/MS; **(B)** PCA score plot built from selected regions of the normalised FT-IR spectra (SNV, baseline removal and Savitzky-Golay second derivative) of acidified stool samples. Samples are classified according to butyrate levels measured by GC/MS: above the 90th percentile (high butyrate, green squares), below the 10th percentile (low butyrate, orange triangles), or intermediate levels (grey crosses).

### Sample mix and background test

To determine the reliability of the analysis, four samples used in the previous analysis were randomly chosen and mixed in pairs (Supplementary Figure S5). The variance was explained by five PCs, with PC1 and PC2 accounting for 46.15% and 27.03%, respectively (a total of 73.18%). The mixed samples clustered close to or between their respective original samples, confirming the consistency of the method. As a background control, the acidified water used for the preparation of the supernatants was analysed (Supplementary Figure S6). Its SNV-corrected spectra with baseline removal differed markedly from those of stool samples, further validating the specificity of the analysis.

## DISCUSSION

Short-chain fatty acids (SCFAs) are key metabolites produced by the gut microbiota, providing insight into microbial functionality and stool sample quality for FMT and microbiome research. While GC/MS remains the reference method for SCFA quantification, it is costly, time-consuming, and sensitive to pre-treatment variability [12]. We evaluated Fourier-transform infrared (FT-IR) spectroscopy as a faster, simpler alternative.

In *Clostridioides difficile* infection (CDI), dysbiosis leads to reduced protective metabolites such as SCFAs [28]. This was reflected in FT-IR spectra, with CDI samples showing lower absorbance than healthy donors (Figure 1B) in critical SCFA absorption regions. SCFA-specific spectral regions in PCA enabled full separation of eubiotic and dysbiotic samples (Figure 1A), while full-spectrum analysis also distinguished groups but likely included other metabolite signals (Supplementary Figure 2).

In the sample-mixing experiments, PCA analysis revealed that the mixed samples clustered in proximity to their source samples, reflecting their intermediate spectral characteristics (Supplementary Figure 5), in addition, negative controls confirmed signal specificity (Supplementary Figure 6). On the other hand, butyrate showed the best discrimination, consistent with its greater depletion in CDI compared to acetate or propionate [28,29]. However, FT-IR could not provide accurate SCFA quantification (Figure 2), as overlapping peaks from similar compounds limit specificity, unlike GC/MS, which separates and identifies compounds with high precision. The IR Biotyper®, although not designed for stool samples, is already used in hospital microbial typing [17,18] and could enable rapid, low-cost stool quality screening in clinical settings.

This study has several limitations. FT-IR spectroscopy, as applied here, cannot accurately quantify SCFA levels or discriminate between SCFA isomers because of peak overlap from structurally similar compounds. The instrument used was not optimized for stool analysis, and while IR-related techniques such as ATR-FTIR may improve performance, they were not tested. The study population was limited to CDI patients and healthy donors, representing two extremes of microbiota health, which may not reflect intermediate dysbiosis profiles.

Despite these limitations, FT-IR spectroscopy offers a rapid, cost-effective and minimal-preparation approach for detecting SCFA-related spectral differences in stool samples. Although it cannot replace GC/MS for precise quantification, it may serve as a practical tool for quick metabolic quality screening of FMT material or microbiota samples in hospital environments. Future research should expand the analysis to other relevant microbial metabolites and include patient groups with varying degrees of microbiota disruption.

## ACKNOWLEDGMENTS

We acknowledge the Departments of Microbiology and Biochemistry and Molecular Genetics of Hospital Clínic de Barcelona for their contribution.

## DATA AVAILABILITY

The code required to reproduce the analysis presented in this paper is available in the following GitHub repository: [https://github.com/asayol/sayol-altarriba_et_al_2025]. OPUS binary files available upon request.

## AUTORSHIP & CONTRIBUTION

*Writing-Original Draft*: A.S-A., *Writing-Review & Editing*: J.L.V-C., C.C-P., *Conceptualization*: C.C-P., J.L.V-C., *Methodology:* C.P., I.R., G.C, C.C-P., *Investigation*: A.S-A., A.A., A.V., R.A., J.F., *Formal Analysis:* A.S-A., E.L-M., S.M., J.L.V-C., *Visualization*: A.S-A., E.L-M., *Supervision*: C.P., I.R., G.C, S.M., J.L.V-C., C.C-P.

