## SUPPLEMENTARY FIGURES for "Rapid quantification of short-chain fatty acids by Fourier-transform infrared spectroscopy for microbiota quality assessment"

**Supplementary Figure S1.**

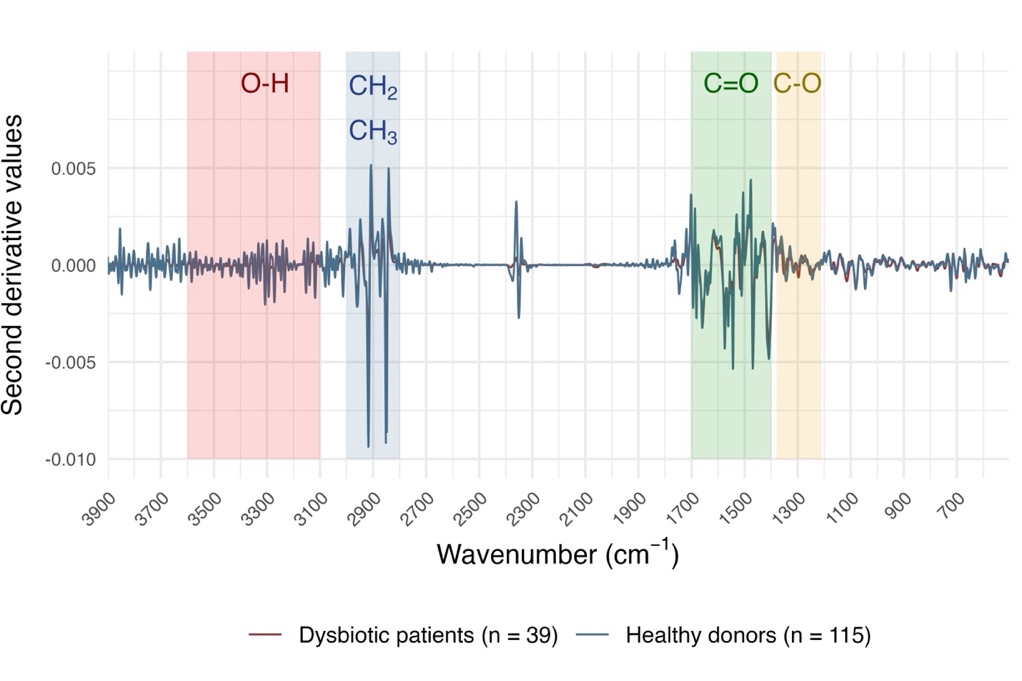

**Supplementary Figure S1.** Second derivative averaged values of mid-infrared (3,992 - 504 cm^-1^) spectra of acidified stool samples from healthy donors (n = 115) (blue) and dysbiotic patients with CDI (n = 39) (red), previously preprocessed with the SNV and the baseline removal. Bands for specific groups and bonds are marked in the spectra: O-H band (3,600 – 3,100 cm^-1^, red), C-H band (3,000 – 2,800 cm^-1^, blue), C=O band (1,700 – 1,400 cm^-1^, green), C-O band (1,380 – 1,210 cm^-1^, yellow).

**Supplementary Figure S2.**

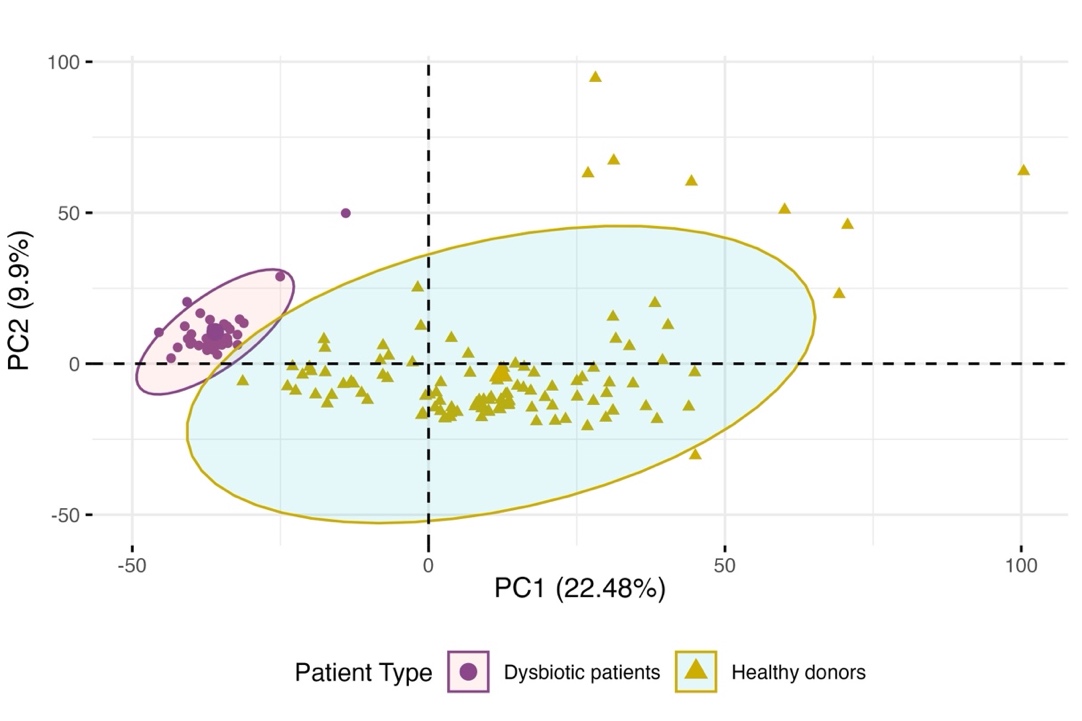

**Supplementary Figure S2.** PCA score plot built from the full normalised FT-IR spectra (3,992 – 504 cm^.1^) (SNV, baseline removal and Savitzky-Golay second derivative) of acidified stool samples. Separation between healthy donors (n = 115) (yellow triangles) and dysbiotic patients with CDI (n = 39) (purple dots), ellipses indicate the 95% confidence interval of each group.

**Supplementary Figure S3.**

| **A)**  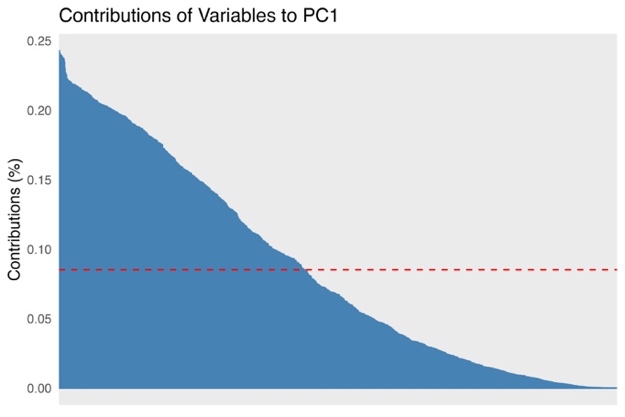 | **B)**  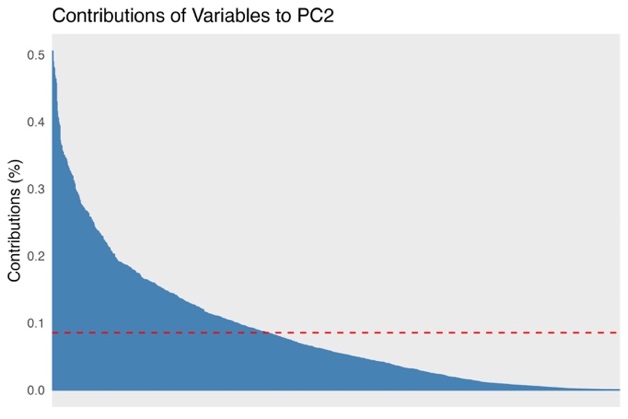 |
| --- | --- |

**Supplementary Figure S3.** Contribution of the 1,174 variables assessed (absorption at selected wavelengths) to the principal components of the PCA: **(A)** PC1; **(B)** PC2. The red dashed line corresponds to the expected value if the contribution were uniform among variable.

**Supplementary Figure S4.**

| **A)**  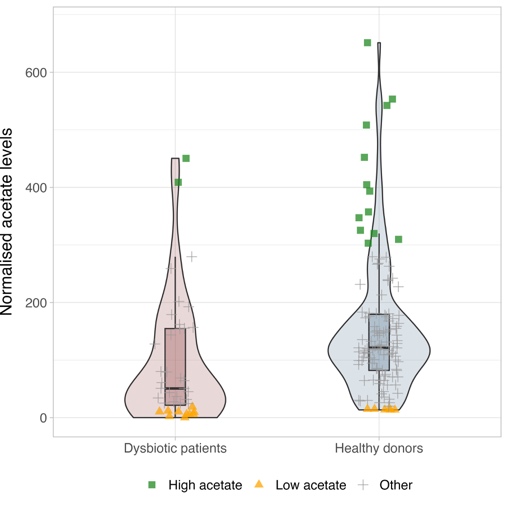 | **B)**  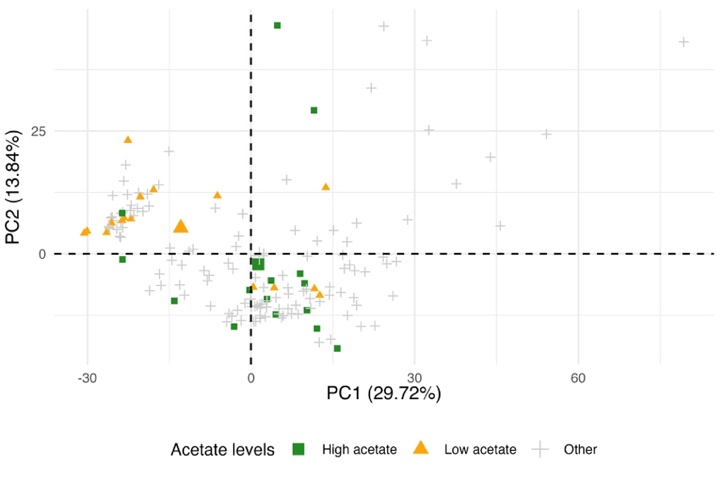 |
| --- | --- |
| **C)**  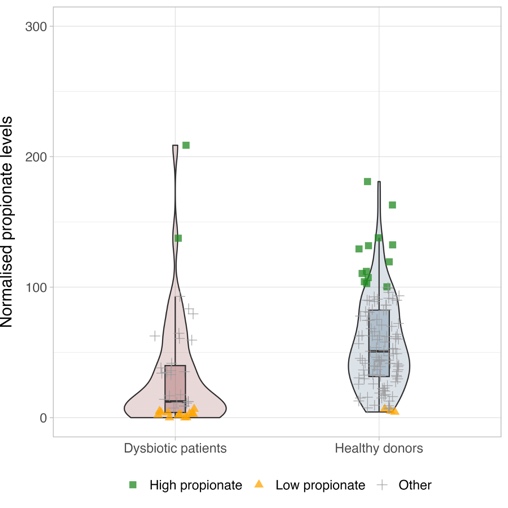 | **D)**  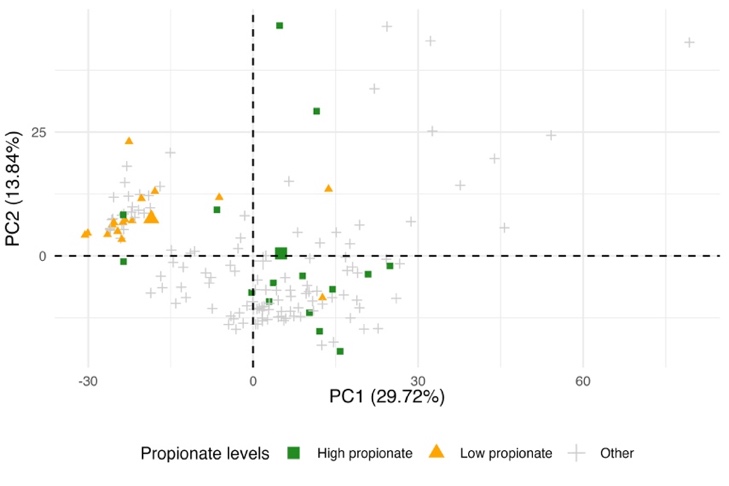 |

**Supplementary Figure S4.** SCFAs levels normalised by bacterial count of dysbiotic patients with CDI (n = 39) (red) and healthy donors (n = 115) (blue) and PCA scoreplot built from selected regions of the normalised FT-IR spectra (SNV, baseline removal and Savitzky-Golay second derivative) of acidified stool samples. Samples are marked according to SCFAs levels above the 90^th^ percentile (green squares), SCFAs levels below the 10^th^ percentile (orange triangles), other levels (gray crosses). **(A)** acetate levels; **(B)** PCA scoreplot of acetate levels; **(C)** propionate levels; **(D)** PCA scoreplot of propionate levels.

**Supplementary Figure S5.**

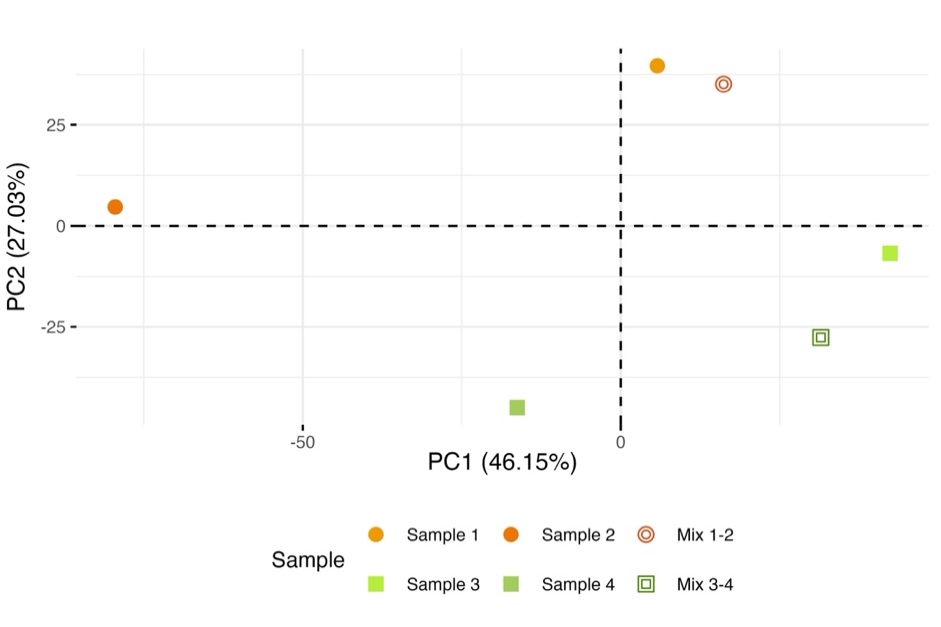

**Supplementary Figure S5.** PCA score plot built from the full normalised FT-IR spectra (3,992 – 504 cm^.1^) (SNV, baseline removal and Savitzky-Golay second derivative) of four random acidified stool samples individually and mixed in pairs. Dots in orange-red: samples 1, 2, and its mix (empty dot); squares in green: samples 3, 4 and its mix (empty square).

**Supplementary Figure S6.**

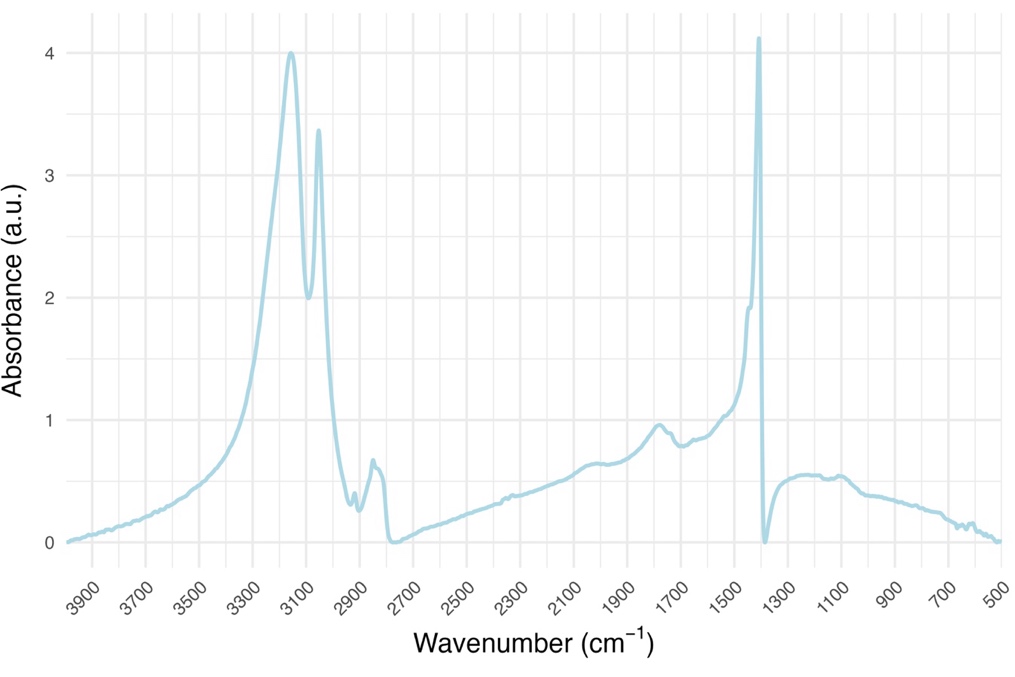

**Supplementary Figure S6.** Absorbance (a.u., arbitrary units) of mid-infrared (3,996 - 500 cm^-1^) spectra of acidified water with HCl.
